# Insights into the Evolution of Ancient Shark and Ray Sex Chromosomes

**DOI:** 10.1101/2025.02.26.637739

**Authors:** Szu-Hsuan Lee, Olivier Fedrigo, Léa Soler-Clavel, Emily Humble, Pierre Lesturgie, Jennifer Balacco, Brian O’Toole, Jacquelyn Mountcastle, Bettina Haase, Nadolina Brajuka, Nivesh Jain, Alan Tracey, Dominic E. Absolon, Sarah Pelan, Damon-Lee Pointon, Jonathan M. D. Wood, Arang Rhie, Daniel J. Macqueen, Kerstin Howe, Eric Jarvis, Gavin J. P. Naylor

## Abstract

While sex-determining mechanisms have been extensively characterized in many vertebrates, they have not been explored in chondrichthyan fishes until relatively recently. In the present study, we used high-quality whole genome reference assemblies to examine the putative sex chromosomes of 14 elasmobranch species spanning nine orders. We describe four newly assembled reference genomes belonging to the white shark *Carcharodon carcharias*, the Atlantic stingray *Hypanus sabinus*, the smalltooth sawfish *Pristis pectinata*, and the zebra shark *Stegostoma tigrinum*. We conducted sex chromosome identification and verification using short-read sequence data collected for multiple individuals for three of the species. This revealed putative pseudoautosomal regions (PARs) and, in one instance, a candidate sex chromosome reassignment. A synteny analysis revealed an ancient and shared origin of the chromosomes within elasmobranchs considerably older than any previously proposed scenario, and a potential candidate gene involved in sex determination shared across all examined species. The synteny analysis also revealed a historical fusion and the formation of neo-Y chromosomes between two myliobatiform species. Our results show that there has been strong conservation and homology of the X chromosomes among elasmobranchs in spite of their varied features and different evolutionary histories.

## Introduction

Of the estimated 75,000 vertebrate species (*IUCN, 2024*), almost all have two sexes. While sex-determination (SD) mechanisms vary across species groups, they are categorized into two systems: Those that are genetically determined (GSD), where the triggers are “hard-wired” and genetic in origin, and those triggered by environmental cues, such as temperature (TSD). The GSD systems are further subdivided into XX/XY systems, where the males are heterogametic, and ZZ/ZW systems, where the females are heterogametic.

The variety of SD systems among jawed vertebrates (Chordata: Gnathostomata) has long intrigued scientists. Most of our understanding is derived from tetrapods (Osteichthyes: Tetrapoda). Therian mammals appear to be mostly XX/XY, birds ZZ/ZW, and snakes both XX/XY and ZZ/ZW, while marine turtles and crocodilians show TSD. Outside of tetrapods, the species-rich and diverse ray-finned fishes (Osteichthyes: Actinopterygii) remain relatively underexplored, but of those examined, many mechanisms have been discovered (Arai, 2011). Members of this group have been shown to exhibit ESD, XY, or ZW systems that are variously homomorphic and heteromorphic, where multiple sex chromosomes are common (Sember et al., 2021), often harboring novel suites of SD genes and mechanisms not seen in other vertebrates (e.g., Bertho et al., 2021; Kikuchi & Hamaguchi, 2013; Pan et al., 2021). Sister to Osteichthyes within the gnathostomes is Chondrichthyes - the cartilaginous fishes. The two groups last shared a common ancestor over 409 million years ago (Miller et al., 2003; Zhu et al., 2009). The chondrichthyans themselves are split into two subgroups: Elasmobranchii, represented by modern sharks, skates, and rays, and Holocephali, or elephant sharks. As of 2024, there are about 1200 valid elasmobranch species (Fricke et al., 2024) and 60 holocephalans.

Until relatively recently, many biological properties of the chondrichthyans were obscured compared to the ray-finned fishes and tetrapods likely due to the difficulty of reliably obtaining high-quality samples. However, once it was discovered that they exhibited a broad range of life history traits, some of which render them vulnerable to extinction, they garnered attention from the conservation community and are now the subject of intense exploration across a variety of disciplines (e.g., Marra et al., 2019; Nakaya et al., 2020; Sendell-Price et al., 2023). In the current study, we set out to greatly expand the growing body of research on chondrichthyans by focusing on the origin and evolution of their sex determination.

The advent of high-throughput sequencing is transforming biology. It is now possible to assemble high-quality reference genomes for species of interest across the Tree of Life (Rhie et al., 2021). These reference genomes allow us to better understand the genes involved in shaping developmental processes, the underpinnings of genetic diseases, and sex determination. Recently, genomic analyses have demonstrated an adoption of XX/XY GSD among some shark and ray species (Wu et al., 2024; Yamaguchi et al., 2023). These studies have contributed to our understanding of the sex determination among Chondrichthyes, as they identified and characterized the sex chromosome pairs, the putative pseudoautosomal regions (PARs), genes, and repeat elements among these sex chromosomes. However, inferences made in these studies have been limited in their taxonomic scope within elasmobranchs.

In this study, we examine the sex chromosomes and their evolution across 14 genomes spanning nine different orders of elasmobranchs and update some previous conclusions based on new evidence. We first examined the assignment of the sex chromosomal pairs in selected species and identified the potential boundaries of PARs. We examined the identified X chromosomes across 13 species across eight elasmobranch orders, which uncovered the consistent presence of *HoxC* genes, as well as a potential candidate for the elasmobranch SD gene, the *ahr*-like (aryl hydrocarbon receptor-like). We characterized four elasmobranch Y chromosomes from three orders and inferred diverse evolutionary trajectories, including chromosome fusions that resulted in neo-Y chromosomes in Myliobatiformes. Lastly, we conducted sex chromosome synteny analyses, which revealed the ancient origin of the elasmobranch sex chromosomes and provided a more comprehensive understanding of the evolution of sex determination within elasmobranchs and, in turn, jawed vertebrates.

## Materials and Methods

### Sample collection

For the reference genome dataset, we collected tissue samples from one individual of each of the following four species: Atlantic stingray (*Hypanus sabinus*), smalltooth sawfish (*Pristis pectinata*), white shark (*Carcharodon carcharias*), and zebra shark (*Stegostoma trigrinum*) from around the United States between 2006 and 2021. Information associated with the tissue samples is summarized in Supplementary Table 1.

Depending on the species, blood, or spleen tissues were collected from the location, transported, and flash frozen in liquid nitrogen.

For the short-read genomic sequencing dataset, fin clips from two male and two female white sharks were collected in South Africa, and fin clips or blood tissues from two male and two female thorny skates were collected in the NorthWest Atlantic (Supplementary Table 1).

### Reference genome assembly and annotation

Chromosome-level reference genomes for the Atlantic stingray, smalltooth sawfish, white shark, and zebra shark were sequenced and assembled by the Vertebrate Genome Lab (Rockefeller University, New York, NY, USA). The assemblies were manually curated, and chromosomal units were identified and named where possible by the Genome Reference Informatics Team (Tree of Life, Sanger Institute, UK).

### The Atlantic stingray genome

Genomic DNA was isolated using the MagAttract HMW DNA Kit (Qiagen 67563). A total of 15 mg of spleen tissue yielded 16 µg of DNA. DNA quantity and quality were assessed with a Qubit 3 fluorometer (Invitrogen, Qubit dsDNA Broad Range Assay, cat no. Q32850) and a Fragment Analyzer (Agilent Technologies, Santa Clara, CA), respectively. Shearing was performed on 10 µg of DNA using a Megaruptor 3 (Diagenode, Denville, NJ, USA), and a Hifi PacBio library was prepared with the PacBio SMRTbell Express Template Prep Kit 2.0. The library was size-selected (>10 kb) on a Pippin HT (Sage Science, Beverly, MA, USA), yielding 1.7 µg of final library material with an average size of ∼15 kb. Sequencing was carried out on six 8M SMRT Cells on the Sequel II platform, with 30-hour movies.

Hi-C sequencing was performed using the Arima v2 Hi-C kit, yielding ∼77x coverage on a NovaSeq 6000 platform.

Genome assembly used the VGP pipeline v2 (Hifiasm + Hi-C phasing v. 0.16.1) on the Galaxy platform, producing contigs from PacBio HiFi long reads, with Hi-C data scaffolding using yahs v. 1.2a (Larivière et al., 2024). The genome assembly was manually curated by the Wellcome Sanger Institute’s Genome Reference Informatics Team under the ID of sHypSab1.

### The smalltooth sawfish genome

Thirty mg of spleen tissue provided 23 µg of DNA following the Bionano Prep Animal Tissue DNA Isolation Fibrous Tissue Protocol (Document Number 30071). DNA quantity and quality were assessed with a Qubit 3 fluorometer and Fragment Analyzer, respectively. Sixteen µg of DNA was sheared using a 26G blunt-end needle following PacBio protocol PN 101-181-000 (Version 05), and a large-insert library was prepared using the PacBio Express Template Prep Kit v2.0 (#100-938-900). This library was size-selected (>20 kb) on a BluePippin Size-Selection System, resulting in a 1.8 µg Continuous Long Read (CLR) library, averaging ∼49 kb in size, and was sequenced on 17 SMRT Cells on the Sequel I platform (10-hour movies).

Bionano Genomics optical mapping was performed on 750 ng of DNA, isolated using the Bionano Prep protocol, quality-checked with pulsed-field gel electrophoresis (Pippin Pulse, Sage Science, Beverly, MA), and labeled using the Bionano Prep Direct Label and Stain (DLS) Protocol (Document Number 30206). The run was performed on a Bionano Saphyr platform

Hi-C sequencing was carried out by Arima Genomics, with ∼74x coverage. Additionally, Linked-reads library preparation was performed using the 10X Genomics Chromium platform (Genome Library Kit & Gel Bead Kit v2 PN-120258, Genome Chip Kit v2 PN-120257, i7 Multiplex Kit PN-120262). The prepared library was sequenced on an Illumina Novaseq S4 150bp PE lane platform at an average 65x coverage.

Genome assembly was performed using the VGP pipeline v1.5. FALCON v. DNANexus 1.9.0 and FALCON-Unzip v. DNANexus 1.0.6 were used to generate contigs from CLR. purge_haplotigs v. v1.0.3 and 1.Nov.2018 were applied to reduce false duplications from the primary contigs. For scaffolding, scaff10x v. 4.1.0 was used to scaffold the 10x data, Bionano Solve DLS v. 3.2.1 for the Bionano data, and Salsa HiC v. 2.2 for the Hi-C data. The genome assembly was manually curated by the Wellcome Sanger Institute’s Genome Reference Informatics Team with gEVAL manual curation v. 2019-11-18 (Howe et al., 2021) under the ID of sPriPec2. Arrow smrtanalysis Pacbio polishing & gap filling v. 6.0.0.47841 were applied to polish the PacBio contigs, while longranger align v. 2.2.2 and freebayes Illumina polishing v. 1.3.1 were applied to benchmark and polish the assembly with Illumina data.

### The white shark genome

DNA was extracted from a blood sample taken from a live animal using the Bionano Prep Animal Tissue DNA Isolation Fibrous Tissue Protocol. The protocol yielded 43 µg of total DNA. A 10 µg sample of total DNA was sheared using a 26G blunt-end needle (Pacbio protocol PN 101-181-000 Version 05). The library was prepared with the PacBio Express Template Prep Kit v2.0 (#100-938-900), size-selected (>20 kb) on a BluePippin System, resulting in a 1.7 µg CLR library. The library was sequenced using seven SMRT Cells on the Sequel I platform (10-hour movies).

Optical mapping was performed with 750 ng of DNA using the Bionano Prep Animal Tissue DNA Isolation Fibrous Tissue Protocol (Document Number 30071) protocol. DNA quality was assessed using pulsed-field gel electrophoresis (Pippin Pulse, SAGE Science, Beverly, MA) and then labeled for Bionano Genomics optical map using the Bionano Prep Direct Label and Stain (DLS) Protocol (Document Number 30206). The run was performed on the Bionano Saphyr platform.

Hi-C sequencing was carried out by Arima Genomics, using the Arima-HiC 2.0 kit at ∼100x coverage. Linked-reads library was prepared using the 10X Genomics Chromium Genome Library Kit & Gel Bead Kit v2 PN-120258, Genome Chip Kit v2 PN-120257, i7 Multiplex Kit PN-120262 kit) and sequenced on an Illumina NovaSeq S4 platform with ∼60x coverage.

Genome assembly was carried out using the VGP pipeline v1.6 (Rhie et al., 2021). FALCON v. DX 2.0.1 and FALCON-Unzip v. DX 1.2.1 were used to generate contigs from CLR. purge_dups v. DX 19-08-10 were applied to reduce false duplications from the primary contigs. For scaffolding, scaff10X v. DX 2.0.3 was used to scaffold the 10x data, Bionano Solve v. DX 3.4.0 for the Bionano data, and Salsa2 HiC v. DX 2.2.0 for the Hi-C data. The genome assembly was manually curated by the Wellcome Sanger Institute’s Genome Reference Informatics Team with the gEVAL manual curation v. 2021-02-05 under the ID of sCarCar2 Arrow polishing and gap filling v. DX 7.0.1.1 were applied to polish the PacBio contigs, while longranger align v. 2.2.2.1 and freebayes DX v. 19-07-11 were applied to benchmark and polish the assembly with Illumina data.

### The Zebra shark genome

A 30 µL whole blood sample was used to extract and isolate 17 µg of DNA with the Nanobind CBB Big DNA Kit (NB-900-001-01) and with the Nanobind UHMW DNA Extraction protocol. DAN quantity and quality assessments were completed with a Qubit 3 fluorometer (Invitrogen Qubit dsDNA Broad Range Assay cat no. Q32850) and Fragment Analyzer (Agilent Technologies, Santa Clara, CA), respectively. Fifteen µg of DNA was sheared with a Megaruptor 3 (Diagenode, Denville, NJ, USA), followed by a HiFi library preparation using the PacBio SMRTbell Express Template Prep Kit 2.0. Size selection on a Pippin HT (Sage Science, Beverly, MA, USA) resulted in a 1.6 µg library with an average insert size of ∼20 kb. Sequencing was performed on four SMRT Cells on the Sequel II platform with 30-hour movies.

Optical mapping used 750 ng of DNA, isolated using the Nanobind CBB Big DNA Kit (NB-900-001-01). DNA quality was assessed using the pulsed-field gel electrophoresis (Pippin Pulse, SAGE Science, Beverly, MA) and labeled using the Bionano Prep Direct Label and Stain (DLS) protocol (document number 30206). The run was performed on a Bionano Saphyr instrument. Hi-C sequencing was conducted on an Arima v2 Hi-C kit, with the library sequenced on a NovaSeq 6000 platform at ∼100x coverage.

The genome assembly was generated using the VGP pipeline v2. Hifiasm + Hi-C phasing v. 0.16.1 on galaxy3 to generate contigs from PacBio HiFi long reads and to scaffold the Hi-C data. Additionally, yahs v. 1.2a.2 on galaxy0 was applied to scaffold the Hi-C data. Bionano Solve v. 3.7.0 on galaxy0 was performed to scaffold the Bionano data. The genome assembly was manually curated by the Wellcome Sanger Institute’s Genome Reference Informatics Team under the ID of sSteFas4.

Annotations of the four genomes were generated using the NCBI Eukaryotic Genome Annotation Pipeline (https://www.ncbi.nlm.nih.gov/refseq/annotation_euk/process/) performed by the National Center for Biotechnology Information (Bethesda, MD, USA).

### Reference genome dataset

The reference genome dataset used herein comprises 14 elasmobranch chromosome-level assembled and annotated genome assemblies collected from three sources: (1) the four assembled and annotated reference genomes of Atlantic stingray, Smalltooth sawfish, White shark, and Zebra shark, generated by the VGP that we herein describe. (2) The reference genomes of the thorny skate (sAmbRad1.1.pri, GCF_010909765.2), epaulette shark (sHemOce1.pat.X.cur., GCF_020745735.1), sharpnose sevengill shark (*Heptranchias perlo*, sHepPer1.hap1, GCF_035084215.1), horn shark (*Heterodontus francisci*, sHetFra1.hap1, GCF_035084215.1), Caribbean electric ray (*Narcine bancroftii*, sNarBan1.hap1, GCF_036971445.1) in our reference genome dataset. These genomes were all assembled by the VGP, and the assembly and annotation methods of the thorny skate and epaulette shark reference genomes were previously described by Rhie et al. (2021) and Sendell-Price et al. (2023), respectively. (3) Five further genomes with NCBI RefSeq annotations from the NCBI. These include Lesser devil ray (*Mobula hypostoma,* GCF_963921235.1), Little skate (*Leucoraja erinacea*, GCF_028641065.1), Smaller spotted catshark (*Scyliorhinus canicula*, GCF_902713615.1), Whale shark (*Rhincodon typus*, GCF_021869965.1), and White-spotted bamboo shark (*Chiloscyllium plagiosum*, GCF_004010195.1). The associated information for each genome included in the dataset is summarized in Table 1. We chose to include elasmobranch genomes with predicted genes that are annotated by RefSeq (https://www.ncbi.nlm.nih.gov/refseq/).

### Short-read resequencing data collection

We leveraged short-read resequencing data for several target species to verify the identity of sex chromosomes and examine PARs. We specifically obtained short-read resequencing data from two male and female individuals of three species: the epaulette shark, thorny skate, and white shark. The thorny skate and white shark tissue samples were stored in the 95% ethanol solution at 4° C until the DNA extraction was performed. Extraction of DNA was conducted using E.Z.N.A. Tissue DNA Kit (Omega Bio-Tek, Inc., Norcross, GA, USA), following the manufacturer’s protocols. Samples were delivered to the UF ICBR (Gainesville, USA) for library preparation using NEBNext Ultra II DNA Library Prep Kit for Illumina (New England Biolabs, Inc., Ipswich, MA, USA), which were sequenced at 20x estimated coverage using an Illumina Novaseq 6000 sequencing platform. The thorny skate short-read sequencing data were previously described in Lesturgie et al. (2025).

In addition, Illumina short-read data for two male and two female epaulette sharks kindly provided by the Schartl Research Group (University of Wüzburg, Wüzburg, Germany) were included in the short-read dataset for this species (Supplementary Table 1).

### Short-read sequences mapping and read coverage analyses

For the Illumina reads, adapter sequences and low-quality regions (<Q20) were trimmed with Trimgalore v0.6.10 using default options (https://github.com/FelixKrueger/TrimGalore). Trimmed reads were mapped to the reference genome assemblies of corresponding species using BWA-MEM v0.7.17 (Li, 2013), followed by marking and removal of duplicate reads with Picard (v3.0.0) MarkDuplicates (https://broadinstitute.github.io/picard/).

Sex chromosome assignment and verification and the identification of putative pseudoautosomal region (PAR) boundaries were determined, where possible, using read coverage ratios. To verify the sex chromosome scaffolds and identify putative PARs, the coverage ratios were calculated on a per base pair basis across the chromosome and then compared to the expectation of ratios across X, Y, and autosomal scaffolds, which are 2:1, 0:1, and 1:1, respectively. This method, known as the Chromosome Quotient (Hall et al., 2013), was adopted in previous elasmobranch studies (Wu et al., 2024; Yamaguchi et al., 2023) for the identification of sex chromosome scaffolds.

As suggested by Palmer et al. (2019), stringent mapping criteria should be applied for coverage-based sex chromosome identification. The homologous genes shared between the X and Y chromosomes, known as gametologs, can retain regions with highly similar sequences that remained recombining until relatively recently. Incorrect mapping of short reads onto these regions can diminish the coverage differences observed between the heterogametic chromosomes, causing false negative inferences (Palmer et al., 2019). Cognizant of these concerns, we implemented a quality filter through SAMtools v1.15 – v1.20 (Danecek et al., 2021), set at the minimal MAPQ = 60, to retain only the reads mapped to the reference at the minimum error rate for BWA-MEM. The reads from the samples of each species were pooled according to sex. Pooled sequences were then standardized using the total post-filtered read counts mapped to the reference genomic sequences (Supplementary Table 2), followed by a calculation of the female-to-male coverage ratios along each target chromosome in 10,000 bp, non-overlapping windows, which we found to perform optimally in downstream analyses and visualization when testing various window sizes.

### Comparisons of genes on the sex chromosomes

Gene searches and comparisons for the assigned sex chromosomes were based on the annotated gene lists according to the NCBI RefSeq annotations. Annotated genes were exported from the corresponding genome assembly pages. A subset of genes with locations on the identified or inferred X chromosomes (Table 2) was extracted for each species, which include the Atlantic stingray (chrX1), epaulette shark, horn shark, lesser devil ray (chrX1), little skate (inferred as chr40, see results), thorny skate (reassigned to chr46), whale shark (labeled as chr41, Yamaguchi et al., 2023), white shark, whitespotted bambooshark (inferred as chr43, see results), sharpnose sevengill shark, smaller spotted catshark (labeled as chr28 in the annotated assembly, sScyCan1.1), smalltooth sawfish, and zebra shark. Genes annotated on the X unlocalized scaffolds --scaffolds that are associated but ambiguously joined with the X chromosome (Rhie et al., 2021) --were retained. The intersection of this X-linked gene subset is examined with the gene names annotated by NCBI. The intersection of the Y-linked genes is analyzed with the same method but includes only the genomes where the Y chromosomes are identified, which were for the Atlantic stingray, epaulette shark, lesser devil ray, and white shark. We specifically targeted the locations of *HoxC* genes and *zbtb39*, as these two genes were previously identified from the putative XPAR and Y chromosomes of other elasmobranch species (Wu et al., 2024; Yamaguchi et al., 2023).

### Synteny analyses

Chromosomal synteny was assessed using predicted protein sequence data retrieved from repositories of the 14 genome assemblies on the NCBI (https://www.ncbi.nlm.nih.gov/) and analyzed following the method described by Yamaguchi et al. (2023). Sequences derived from alternative splicing were removed using custom scripts, retaining sequences without isoforms or labeled as isoform X1, while ortholog groups were searched using SonicParanoid v2.0.4 (Cosentino & Iwasaki, 2019) with the ‘-m sensitive’ option. 1-to-multiple and multiple-to-multiple ortholog pairs were then removed, and the conserved synteny based on the 1-to-1 orthologs among species pairs was then searched against the NCBI Entrez database using R package rentrez v1.2.3 (Winter, 2017) to retrieve their chromosomal positional information and then plotted using RIdeogram (Hao et al., 2020).

We assessed the number and identities of X-linked protein-coding genes conserved across the 13 reference-genome assemblies where X chromosomes were assigned or reliably inferred. This excluded the Caribbean electric ray genome since we cannot unambiguously infer the X chromosome due to the lack of support from coverage differences or the evidence of multiple and unique chromosomal fusions involving the syntenic block used to infer the X chromosomes in other species (see Results).

Conserved X-linked genes among the remaining 13 genomes from the genomic dataset were assessed using the methods described above, except for data extracted from the 1-to-1 ortholog groups identified from all included species. We note that this provides a conservative estimation of the number of shared X-linked genes. This is because while the focus of the 1-to-1 ortholog groups is to avoid erroneous inferences caused by paralogous copies of genes, the criterion removes any ortholog for which a duplication was detected in *any* of the 13 species examined. This means that if a gene was observed to be single-copy and putatively orthologous in 12 species but duplicated or missing from the chromosome (see Discussion) in a single species, it would be removed from the count.

Pairwise chromosomal synteny among batoid species based on genomic sequences was used to validate the general patterns of observed synteny. A genome-wide sequence alignment of the Atlantic stingray, lesser devil ray, little skate, manta ray (*Mobula birostris*), and thorny skate was performed using Progressive Cactus (Armstrong et al., 2020), a resource that will be described in a separate manuscript. Syntenic relationships were inferred using HalSynteny (Krasheninnikova et al., 2020), with the options –maxAnchorDistance 1000000 –minBlockSize 200000 and the results plotted using RIdeogram (Hao et al., 2020).

## Results

### Genome assemblies produced by the VGP

Metrics of the four previously undescribed genomes assembled by the VGP (Atlantic stingray, smalltooth sawfish, white shark, zebra shark) are summarized in Fig. 1 and Table 1.

**Figure 1.**
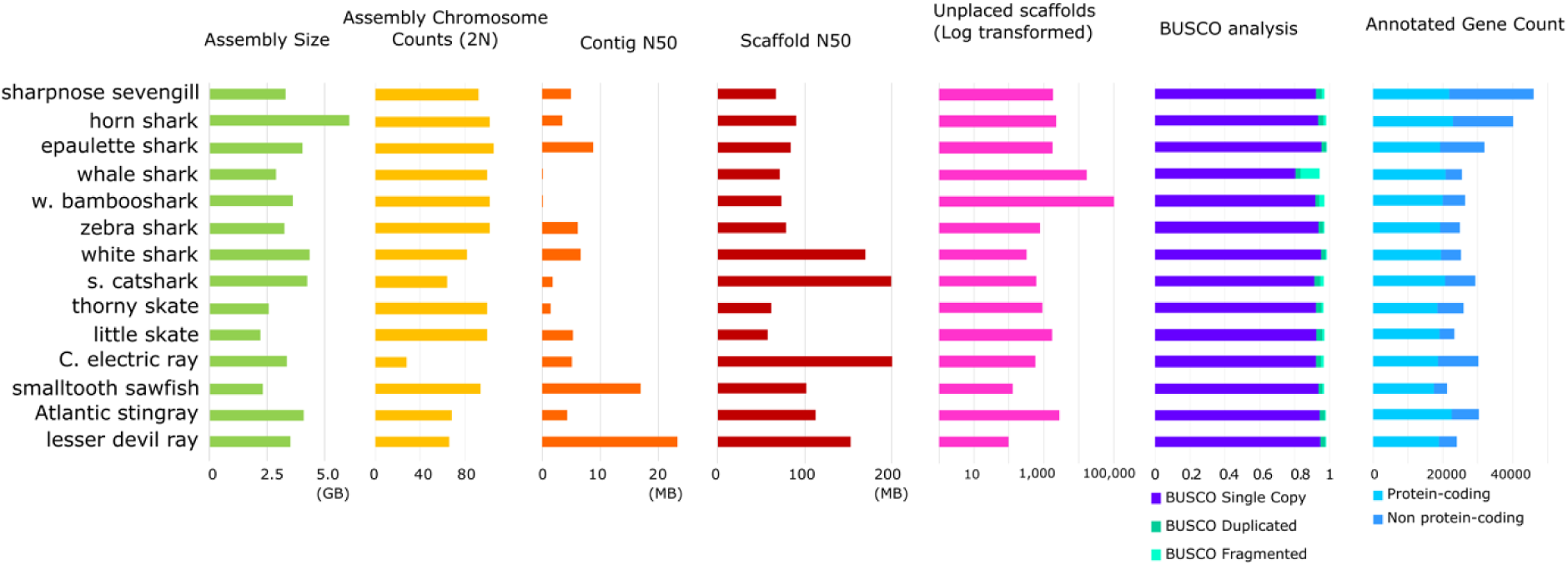
Statistics of the annotated reference elasmobranch genome assemblies examined.

The Atlantic stingray genome assembly is 4 Gb with 32 autosomes, 2 X chromosomes (denoted as X1 and X2), and a Y chromosome. 4,637 contigs were resolved into 2989 scaffolds of which 35 were identified as chromosome level and another 137 unlocalized, i.e., with a known association with a chromosome but ambiguous placement within, representing 89.8% of the assembly sequence. A further 2817 scaffolds (10.2% of the assembled sequence) remain unplaced (Supplementary Fig. 1). The scaffold N50 is 112.3 Mb, the scaffold L50 12, and the contig N50 4.3 Mb. A total of 30,308 genes were annotated, 22,522 of which were protein-coding. Based on the conserved vertebrate gene set, vertebrata_odb10 lineage dataset (n = 3,354), the genome showed 97.1% BUSCO completeness (Table 1). 51.22% of the genome sequences were masked with WindowMasker (Morgulis et al., 2006).

The smalltooth sawfish genome assembly is 2.3 Gb with 46 autosomes and an X chromosome. 870 contigs were resolved into 172 scaffolds, of which 46 chromosomes were identified as chromosomal level, representing 99.6% of sequences. A further 126 scaffolds (0.4%) remain unplaced (Supplementary Fig. 1). The scaffold N50 is 101.7 Mb, the scaffold L50 10, and the contig N50 17 Mb. A total of 21,162 genes were annotated, 17,434 of which were protein-coding. BUSCO completeness was 96% (Table 1) and 37.67% of the genome was masked.

The white shark genome assembly is 4.3 Gb with 41 autosomes, an X chromosome, and a Y chromosome. 3,357 contigs were resolved into 718 scaffolds, of which 42 were identified as chromosome level and another 361 unlocalized, representing 99.6% of the assembly sequence. A further 315 unplaced scaffolds (0.4%) remain unplaced (Supplementary Fig. 1). The scaffold N50 is 169.9 Mb, the scaffold L50 11, and the contig N50 6.5 Mb. A total of 25,164 genes were annotated, 19,440 of which were protein-coding, BUSCO completeness was 97.7% (Table 1) and 51.57% of the genome was masked.

The zebra shark genome assembly is 3.2 Gb, comprising 50 autosomes and an X chromosome. 1,994 contigs were resolved into 835 scaffolds, of which 42 were identified as chromosome level 51 chromosomes and another 13 unlocalized, representing 93.4% of sequences. A further 771 scaffolds (6.6%) remain unplaced (Supplementary Fig. 1). The scaffold N50 is 79 Mb, the scaffold L50 14, and the contig N50 6.2 Mb. A total of 24,913 genes were annotated, 19,223 of which were protein-coding. BUSCO completeness was 96.2 % (Table 1) and 41.90% of the genome was masked.

### Summary of the sex chromosomes identified in reference genomes

Statistics of the sex chromosomes for the 14 species are summarized in Table 2. No species were recognized as having a ZZ/ZW system or environmental sex determination. 11 had X chromosomes identified during the assembly or annotation processes (Table 2), while two species (Atlantic stingray and lesser devil ray) were discovered to have a second X chromosome.

Four species (Atlantic stingray, lesser devil ray, white shark, and zebra shark) were identified to have a Y chromosome. The annotated, identified (thorny skate), or inferred (white-spotted bamboo shark and little skate, Table 2) X chromosomes have higher gene densities relative to their autosomes than observed for the chicken Z and human X chromosomes (Bellott et al., 2010). The Y chromosomes are gene-rich in the lesser devil ray and Atlantic stingray but gene-poor in the epaulette shark and white shark.

The two myliobatiform species, lesser devil ray and Atlantic stingray, are unique among the genomes examined in that they possess multiple sex chromosomes, X1X1X2X2/X1X2Y. The X1 and X2 chromosomes of the two myliobatiform genomes (45.0 Mb – 73.0 Mb) are bigger than the X chromosomes of other species examined (16.4 Mb – 28.8 MB) and contain more genes (Table 2).

### Sex chromosome assignment verification

The results of the sex chromosome assignment validation are presented in Fig. 2. Both the X and Y chromosomes were identified in the epaulette shark and white shark genome, while only the X chromosomes were found in the thorny skate genome. The female-to-male coverage ratios of the epaulette shark (Fig. 2a, b) and the white shark (Fig. 2e, f) support the assigned identity of the annotated X and Y chromosomes when compared to coverage observed in the autosomes (Fig. 2g-i, Supplementary Fig. 2). In both cases, the f-to-m short reads coverage ratio on the autosome, X chromosome, and Y chromosome, are close to the expected values of 1, 2, and 0, respectively. However, the labeled X chromosome of the thorny skate reference genome showed a female-to-male coverage ratio close to 1 (Fig. 2c), as would be expected for an autosome (Supplementary Fig. 2), while the previously labeled chr46, was found to exhibit the ratio of 2, as expected in the X chromosome (Fig. 2d). While we cannot identify the Y chromosome in the thorny skate, the evidence based on coverage suggests a reassignment of thorny skate X to the chr46 of the genome assembly.

**Figure 2.**
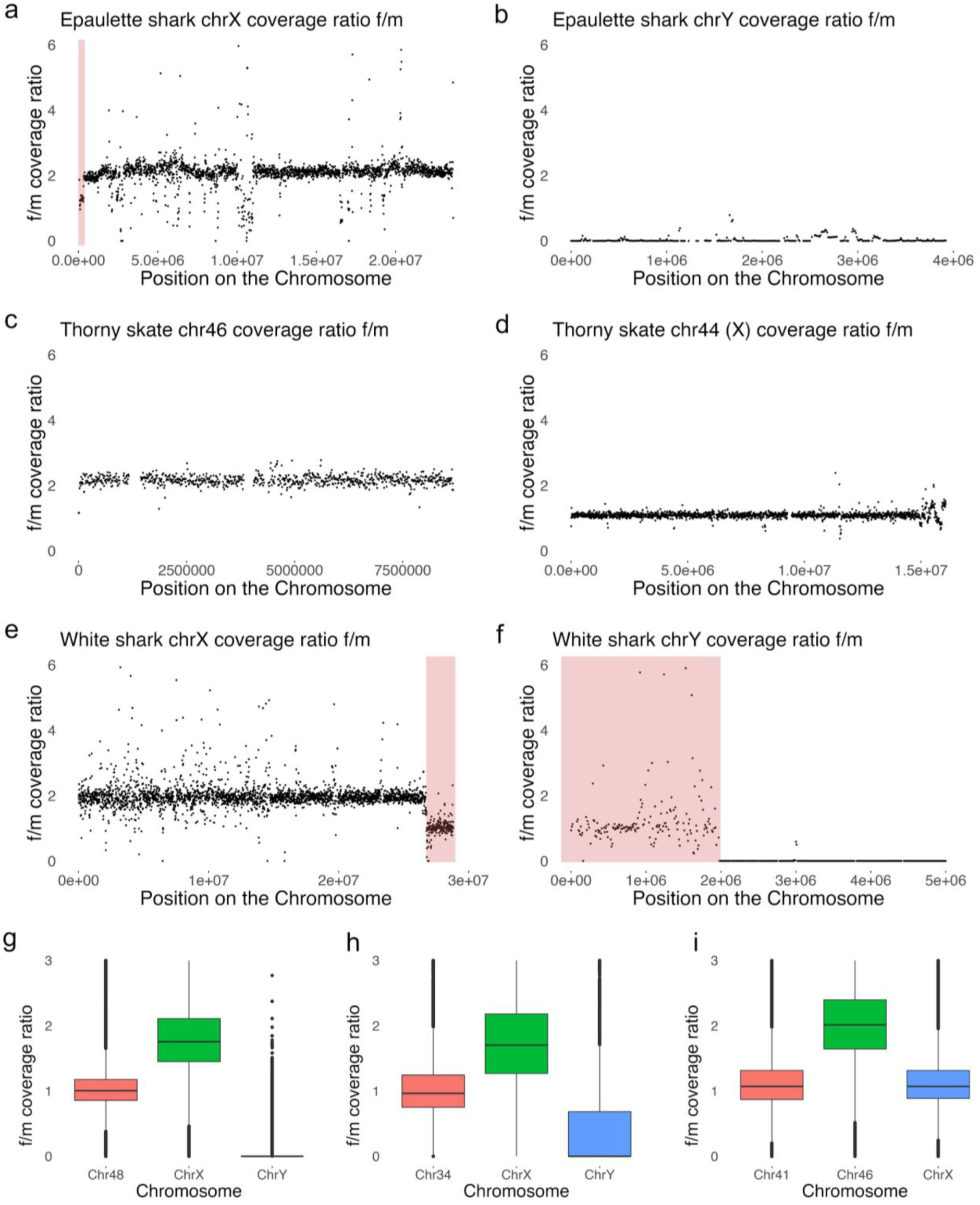
Pooled female-to-male read coverage ratio across sex chromosomes based on the mapping of short-read sequencing data to the RefSeq annotated genome assemblies. Ratios along the chromosomes are visualized in 10,000 bp, non-overlapping windows. (a, b) The f-to-m mapping coverage ratio across chrX and chrY of the epaulette shark genome. (c, d) The f-to-m mapping coverage ratio across chrX and chrY of the white shark genome. (e, f) The f-to-m mapping coverage ratio across chrX and chrY of the white shark genome. Inferred ranges of PARs were highlighted in light red. (g, h, i) Boxplots that show the female-to-male sequencing coverage of identified sex chromosomes and an autosome of the epaulette shark (g), white shark (h), and thorny skate (i).

As shown in previous studies (Wu et al., 2024; Yamaguchi et al., 2023), the female-to-male coverage ratio across sex chromosomes allows the characterization of the boundaries of putative pseudoautosomal regions (PARs). In our results, locations of putative PARs are suggested based on the differences in coverage seen on the X chromosomes of the epaulette shark and white shark and the Y chromosome of the white shark (Fig. 2). Our data support a ∼2Mb PAR at the terminus of the X and the start of the Y chromosome of the white shark genome (but see Discussion). At the start of the epaulette shark X chromosome, a ∼300 Kb long region shows a decrease in female-to-male coverage ratio when compared to the rest of the chromosome, which may represent a putative PAR, although it appears fragmented and has no corresponding region from the Y chromosome. Fragments of the putative PARs are suggested by the coverage ratio on unlocalized scaffolds (Supplementary Fig. 3) associated with the X and Y chromosomes. We found no evidence of a PAR on the thorny skate X chromosome.

### Genes associated with the sex chromosomes

We examined the identity and distribution of genes on both the X and Y chromosomes across species and did not identify a likely master-sex determining (MSD) genes among the ‘usual suspects’ (Herpin & Schartl, 2015; Pan et al., 2021) on the Y chromosome (Table 3). No shared genes were found among the Y chromosomes of the four species where Y scaffolds were identified (i.e., Atlantic stingray, epaulette shark, lesser devil ray, and white shark), and no XY gene pairs were found to be shared across all four species where a Y chromosome was assigned. However, our searches based on the annotated gene names revealed that five genes – *ahr*-like, *igfpb5*-like, *krt8*-like, *nramp2*-like (*slc11a2*-like), and *sp5*-like – were found on the X chromosome across all 13 species. Among which, *ahr* (encoding aryl hydrocarbon receptor), a transcription factor, stands out due to its importance to female and male reproduction in mammals (Baba et al., 2005; Hansen et al., 2014; Pocar et al., 2005) and their interactions with the transforming growth factor-beta (TGF-**β**) signaling pathway (Bussmann & Barañao, 2008; Silginer et al., 2016), a pathway that is important to ovarian and testis development in mammals (Itman et al., 2006; Knight & Glister, 2006) and recurrently gave rise to MSD genes among the teleost fishes (Oliveira et al., 2023; Pan et al., 2021). This *ahr*-like gene, therefore, could be a candidate for a conserved MSD gene among the sharks and rays.

Although we found no putative MSD genes on the Y chromosomes, the *HoxC* cluster was found on the Y chromosomes of the white shark and epaulette shark (Fig. 3; Supplementary Fig. 3).

**Figure 3.**
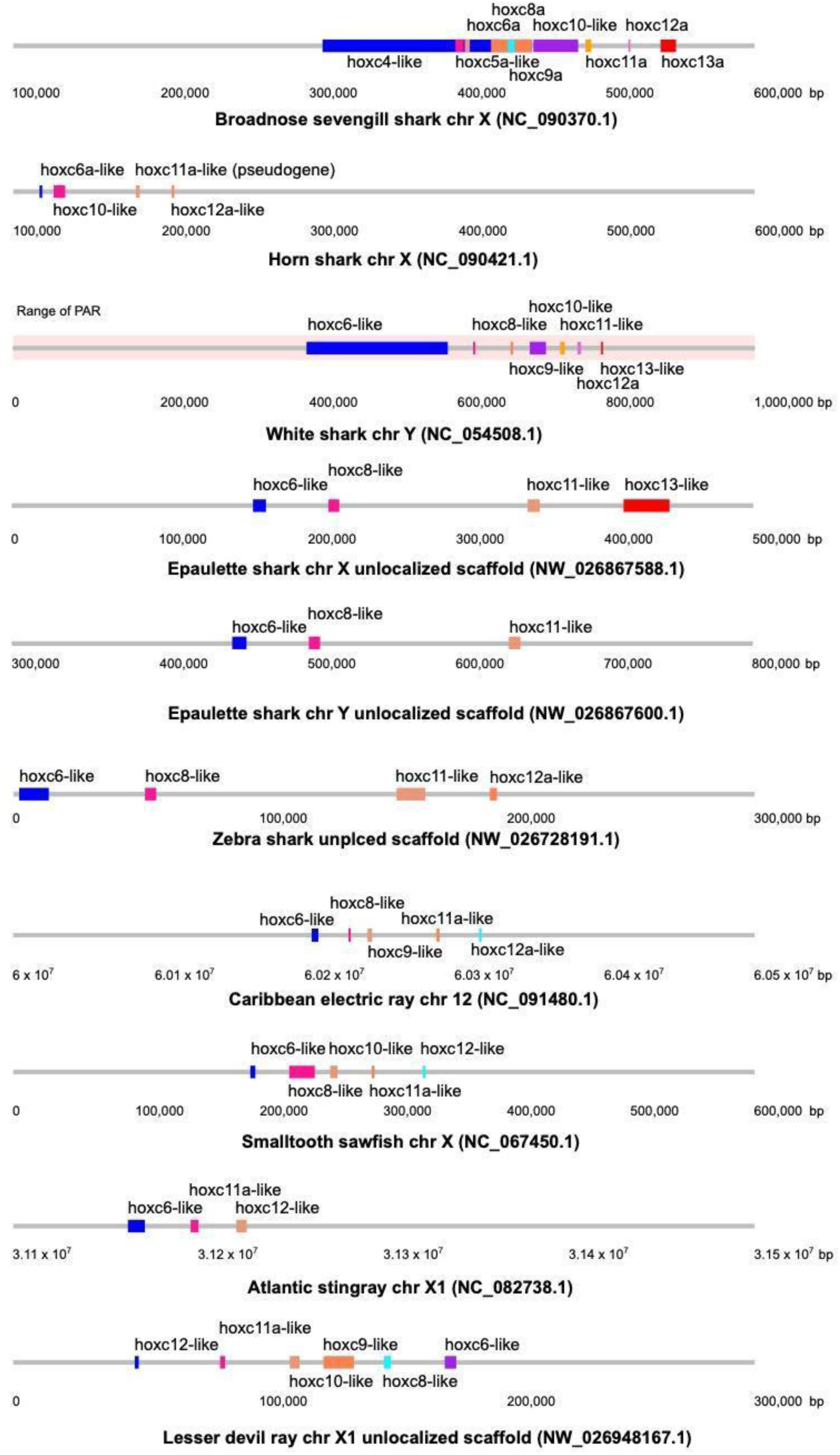
*HoxC* genes in the elasmobranch genomes examined in this study. Note that on the white shark chrY, the *HoxC* genes sit within the putative PAR.

Once thought to be absent among the elasmobranchs (King et al., 2011), the *HoxC* cluster was identified in the brown-banded bamboo shark (*Chiloscyllium punctatum*), whale shark, cloudy catshark (*Scyliorhinus torazame*) (Hara et al., 2018) and found on the XPAR of a zebra shark genome (sSteFas1.1, GCA_022316705.1; Yamaguchi et al., 2023). In the epaulette shark, we found four *HoxC* genes (*hox-c6*, *-c8*, *-c11*, and *-c13*) annotated in one unlocalized scaffold of chrX (NW_026867588.1), while three (*hox-c6, -c8*, and *-c11*) were identified on an unlocalized scaffold of chrY (NW_026728191.1), respectively. By contrast, in the white shark, the Y chromosome harbors seven *HoxC* genes (*hox*-*c6, -c8, -c9, -c10, -c11a, -c12a,* and *-c13*), while none are found on the X chromosome, potentially due to incomplete assembly of the scaffolds (see Discussion). Among these, *hox-c6*, *hox-c8*, and *hox-c11* are shared between the Y chromosomes of the white shark and epaulette shark and are the only Y-linked genes that were found shared between the two species.

*HoxC* genes were not identified in the Y chromosomes of the Atlantic stingray and lesser devil ray, although three (*hox-c6, -c11a,* and *-c12*) and six copies (*hox*-*c6, -c8, -c9, -c10, -c11a,* and *-c12*) are present on the X1 chromosomes (the Atlantic stingray) or X1 unlocalized scaffold (the lesser devil ray), respectively. Despite lacking *HoxC* genes, the Y chromosomes of these two ray species harbor more protein-coding genes when compared to shark counterparts (Table 2), with 24 of these genes shared between the two species as a result of likely fusion events (explored further below).

While lacking Y chromosome assignments, *HoxC* genes were also identified on the X chromosomes of the horn shark, sharpnose sevengill shark, and small tooth sawfish genome. In these species, three (*hox-c6, -c8,* and *-c12a*), nine (*hox-c6a, -c8a, -c9a, -c10, -c11a, -c12a,* and *-c13a*), and five genes (*hox-c6, -c8, -c10, -c11a,* and *-c12a*) were identified on the X chromosomes, respectively, with an extra horn shark *hox*-c11 annotated as a pseudogene. In the Caribbean electric ray genome, where no sex chromosomes have been assigned, five *HoxC* genes were found (*hox-c6, -c8, -c9, -c11a,* and *-c12a*) on chr12.

Additionally, homologs of *zbtb39* are also found on all X chromosomes (X1 for Myliobatiformes) and Y chromosomes, except for the epaulette shark chrY. *zbtb39* was previously identified from the white-spotted bambooshark as one of the only non-recombining X-Y gene pairs, and the X and Y gametologs were found from several elasmobranch species (Wu et al., 2024). In our analysis, we found one copy of *zbtb39* or *zbtb39*-like identified on the X chromosome of 13 out of 14 species, while two copies were identified on the whale shark chrX, suggesting a possible duplication. In Atlantic stingray, lesser devil ray, and white shark, one Y gametolog per species was also identified. Homologs of *zbtb39* were further found on the unplaced scaffolds in epaultette shark (NW_026869144.1), thorny skate (NW_022630193.1), zebra shark (NW_026728070.1). As these three genomes were sourced from male individuals, it is likely that these unplaced scaffolds are associated with the Y chromosomes of the species.

### Comparisons of elasmobranch sex chromosomes via synteny analyses

Our analyses revealed shared synteny among the identified X chromosomes spanning a deep history of elasmobranch evolution (Fig. 4). The pairwise comparison of 1-to-1 ortholog groups indicates that a significant fraction of protein-coding genes located on elasmobranch X chromosomes are conserved across different orders (Supplementary Table 3). Based on the conserved synteny, we identified the putative X chromosomes of both the white-spotted bamboo shark genome assembly (ASM401019v1) and the little skate genome assembly (Leri_hhj_1), as chr43 and chr40, respectively (Supplementary Fig. 4).

**Figure 4.**
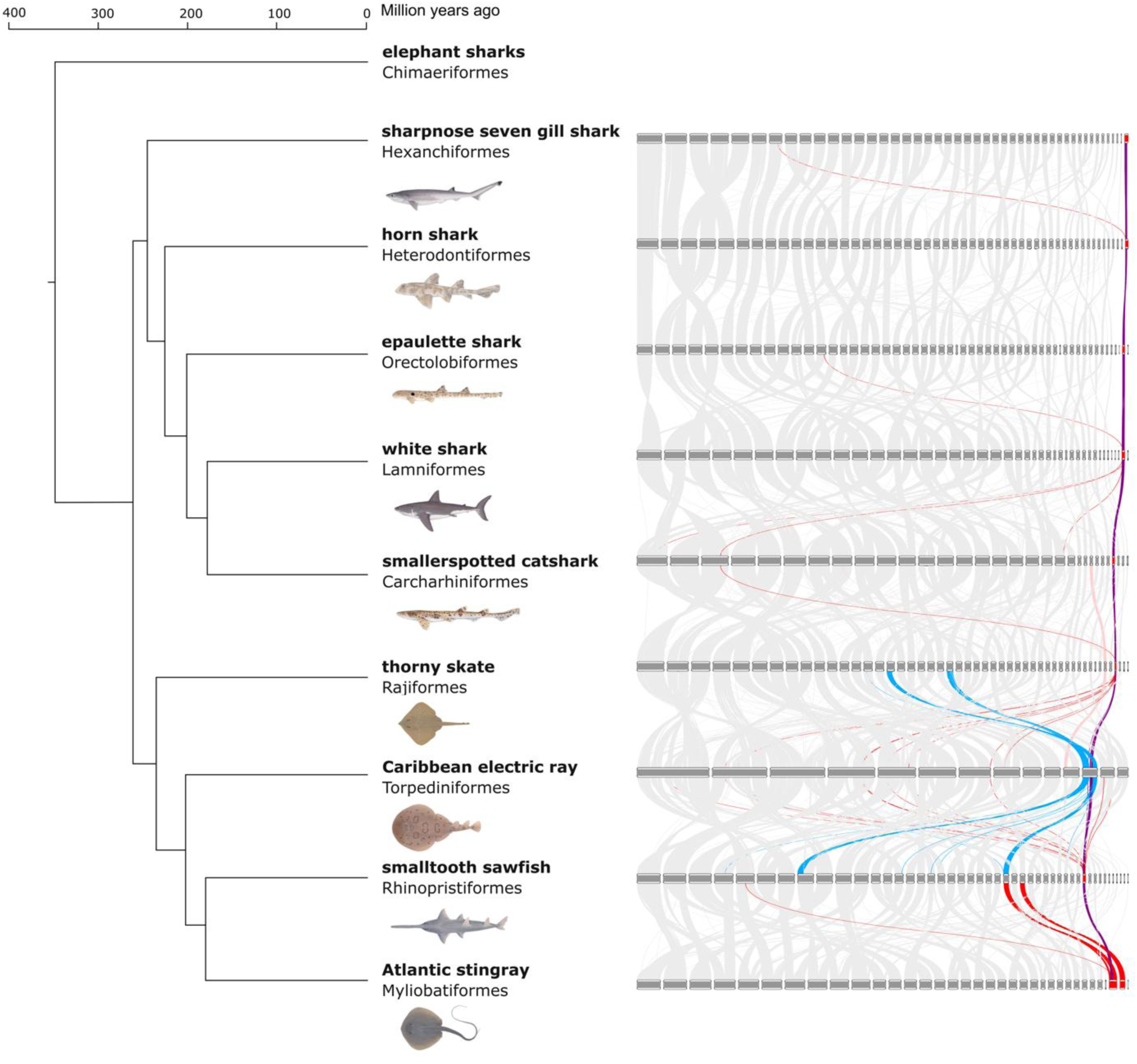
Synteny among the selected species from the six elasmobranch orders, with a focus on the X chromosomes. Syntenic X-linked orthologs between species pairs are highlighted in purple; relationships between orthologs that are X-linked in one species and autosomal in the other are highlighted in red; relationships between orthologs that are potentially X-linked in one species and autosomal in the other are highlighted in blue. Phylogenetic relationships and divergence time estimates are based on Stein et al. (2018).

Homology was also observed between chr12 of the Caribbean electric ray and the conserved X chromosomes of other elasmobranch genomes (Fig. 4), suggesting that this chromosome may be a candidate for the X chromosome of this species (or more broadly for the Torpediniformes). However, this assertion requires further validation, as we identified that this chromosome has undergone several chromosome rearrangements, and clear coverage-based evidence is lacking. Therefore, we excluded the Caribbean electric ray genome in the subsequent analysis of conserved X-linked genes.

Our synteny analysis of the remaining 13 genome assemblies revealed 29 1-to-1 ortholog groups that are conserved among X chromosomes (X1 for the myliobatiform species) across all species (Supplementary Table 4), which we herein described as the elasmobranch X-conserved genes (EXCGs). Of the five genes identified on the X chromosome of all 13 species through the gene search, only three (*ahr*-like, *krt8*-like, and *sp5*-like) were retrieved as ortholog groups among the EXCGs, likely due to the existence of highly similar paralogous copies for the other two genes resulting in their exclusion from our strict definition of 1-to-1 orthologs. These EXCGs are not localized to a single region but are distributed along the entire X chromosome (Fig 5, Supplementary Table 4). These results are consistent with a shared ancestry of elasmobranch X chromosomes, dating back to 265.1 – 348.9 mya (after Stein et al., 2018).

**Figure 5.**
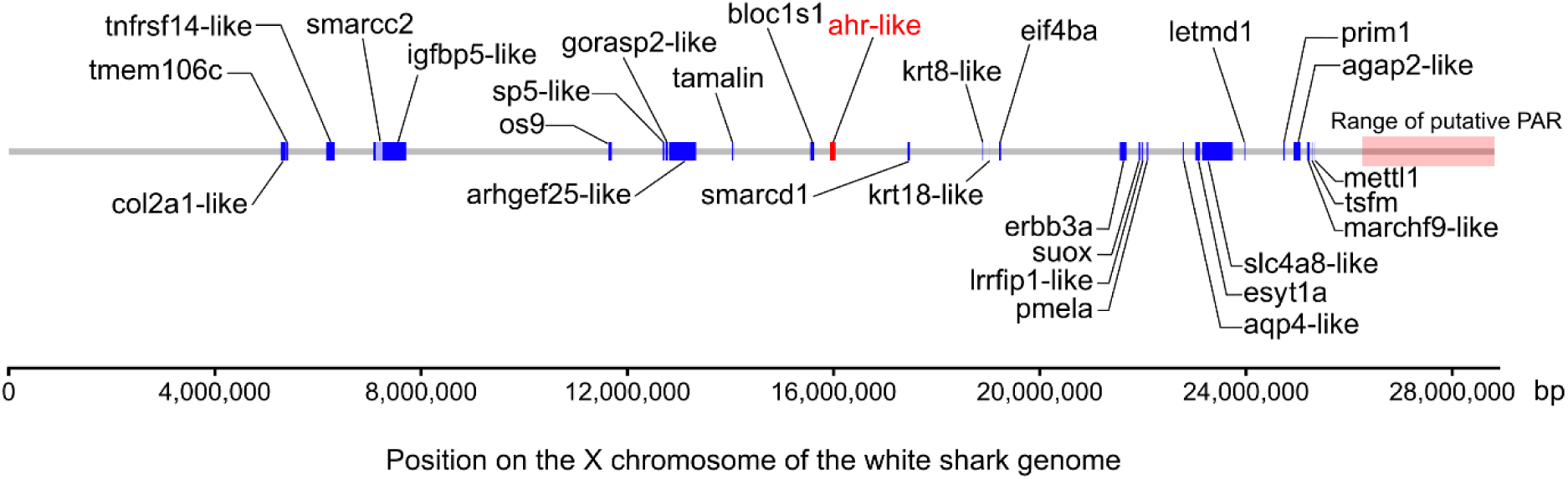
Location of the 29 elasmobranch X conserved genes (EXCGs) on the X chromosome of the white shark reference genome. Presented genes are labeled as their predicted names based on the annotations on the white shark genome. The range of putative PAR and the candidate elasmobranch sex-determining gene, *ahr*-like, are highlighted.

Pairwise synteny analyses further revealed the independent identities of the X1 and X2 chromosomes of the two myliobatiform species, as the two identified X chromosome pairs do not share any syntenic genes (Fig. 6). A block of 138 genes was found to be shared between chrX1 (12,982,165 – 36,032,861 bp) of the Atlantic stingray genome and chrX of the sawfish genome, while 108 genes were shared between chrX1 (11,519,927 – 28,943,168) of the lesser devil ray genome and chrX the sawfish genome. Surrounding these blocks, genes on the remaining chrX1 of the two species are mostly syntenic with those of the sawfish chr25, while chrX2s are syntenic with chr27 of the sawfish reference genome (Fig. 6).

**Figure 6.**
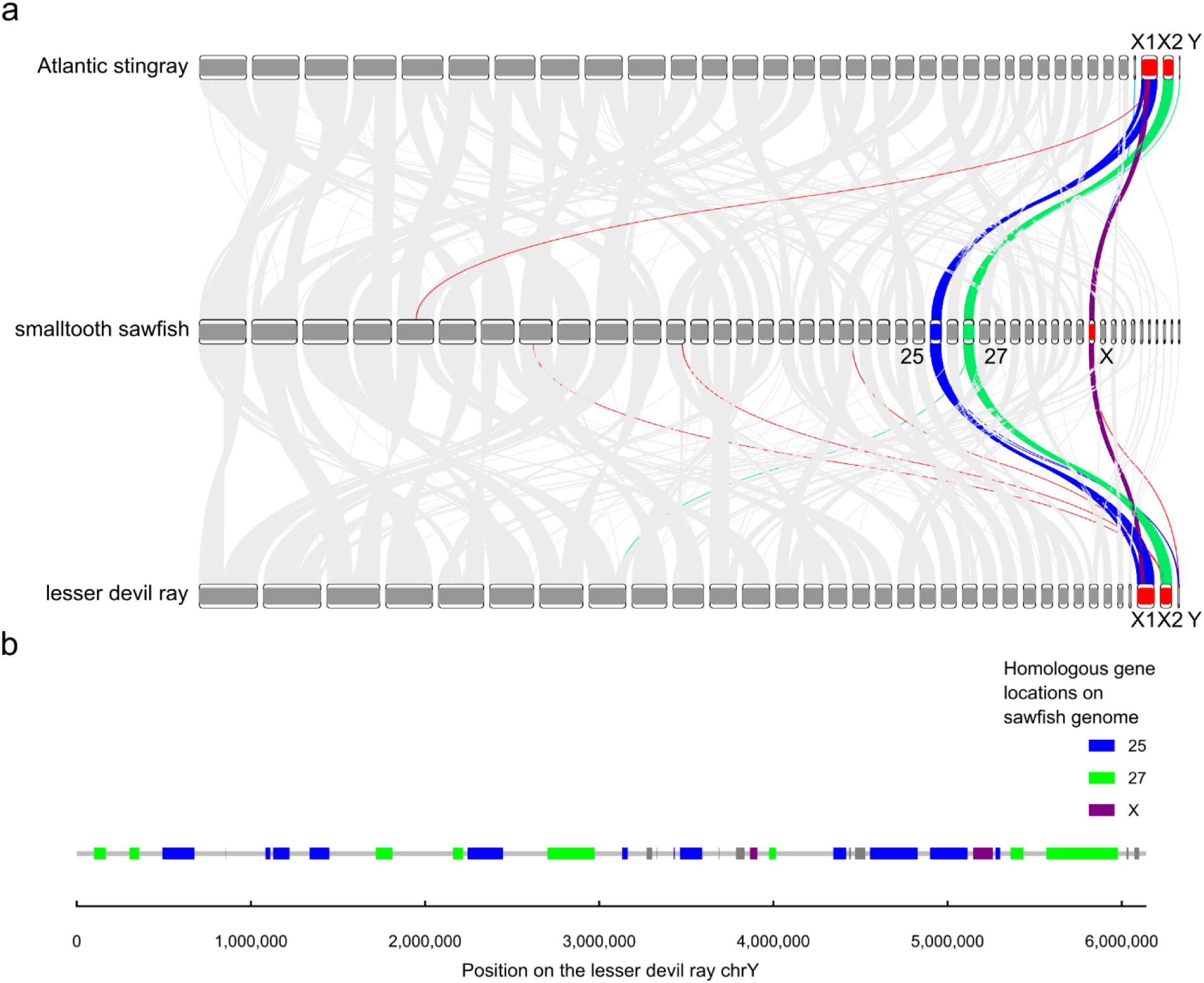
Sex chromosome synteny conservation between the two myliobatiform reference genomes and the smalltooth sawfish genome. (a) Synteny of the sex chromosomes of lesser devil ray, Atlantic stingray, and smalltooth sawfish. (b) Annotated genes on the lesser devil ray chrY, labeled with their homologous location on the smalltooth sawfish genome. Inferred homologous genes and relationships are highlighted in blue if their homologs are located on chr25 of the smalltooth sawfish genome; green, on chr27; purple, on chrX; and red or grey, on other chromosomes.

Interestingly, among the protein-coding genes on the Y chromosome of the Atlantic stingray genome (38 genes) and lesser devil ray genome (40 genes), homologs of 18 genes were found on sawfish chr25, 13 on chr27, and three on chrX (Table 4). These combined results suggest that two chromosomal fusion events occurred after the myliobatiform - rhinopristisform divergence (∼188.0 Mya, Stein et al., 2018). One fusion occurred between the ancestral sex chromosome pair with an autosomal pair (represented by chr25 of sawfish), where X and Y each fused with one autosome homolog. Inversions likely occurred subsequently, shifting the location of the syntenic block to the ancestral X chromosome away from the terminus. The other fusion happened between the Y chromosomes with one homolog of another autosome (chr27 of sawfish) forming a neo-Y chromosome, leaving the unpaired homolog as chrX2 in the two myliobatiform species. This observation is consistent with the proposition made to the X1X2Y system in other myliobatiform species *Potamotrygon falkneri* and *Potamotrygon* aff. *motoro* (Cruz et al., 2011).

Y-linked genes with homologs located on either chr25 or chr27 of sawfish exhibited no regionalization, an observation that is consistent with a history of sequential inversions resulting in recombination arrest (Lahn & Page, 1999) after the proposed fusion events. Although the resulting neo-Y chromosomes in the two myliobatiform species are more gene-rich, they do not seem larger than the Y chromosomes of the species for which fusion events were not inferred (Table 2). This may be the result of either degradation of Y chromosome sequences, as observed across vertebrates including one shark species (e.g., Ezaz et al., 2006; Graves & Peichel, 2010; Wu et al., 2024) and potentially facilitated by the inferred fusions (Pennell et al., 2015), or missing components from the Y chromosomes in the reference genome assemblies.

Chromosomal fusions involving the ancestral X-linked block were also found in another batoid species, the Caribbean electric ray *Narcine bancroftii* (Order Torpediniformes). The genome of this species exhibits the lowest number of chromosomes (2N = 28) among all examined elasmobranch genomes assembled to date, consistent with the karyotype reported for its congener, *Narcine brasiliensis* (Donahue C.S.C., 1974). Our synteny analyses suggest that extensive chromosomal fusions have occurred since the species diverged from the shared common ancestors with the thorny skate (Rajiformes) or the smalltooth sawfish (Rhinopristiformes) (Fig. 4), estimated at 212.0 mya and 209.0 mya (Stein et al., 2018), respectively. These chromosomal fusions, as in the two myliobatiform genomes, included one between the ancestral X-linked block with an autosome homologous to chr25 of sawfish, resulting in the chrX1s observed in Atlantic stingray and lesser devil ray and chr12 of the Caribbean electric ray. However, for electric ray chr12, an additional round of fusion was inferred to have occurred, involving a segment of another autosome (chr8 in sawfish), forming an even larger chromosome (103,180,129 bp). The X2 chromosomes of the myliobatiform genomes (homologous to chr27 of the sawfish genome), are, in turn, homologous to a region on electric ray chr8 as the result of inferred independent fusion events.

## Discussion

### The ancient origin of the elasmobranch sex chromosomes and the putative MSD gene

Our results demonstrate the powerful insights made possible through comparative analyses of high-quality reference genomes that span the evolutionary diversity of extant sharks and rays. Through the analyses of the orthologous genes conserved across the X chromosomes from eight elasmobranch orders, we demonstrate the likely ancient origin of elasmobranch sex chromosome pairs and the dynamic evolutionary trajectories that characterize different elasmobranch lineages. Our results suggest one protein-coding gene among the EXCGs, predicted as aryl hydrocarbon receptor (*ahr*), as a putative candidate for a conserved elasmobranch MSD gene given that MSD genes are usually found in the oldest strata of sex chromosomes (Lahn & Page, 1999; Zhou et al., 2014). Although, to our knowledge, *ahr* has not been shown to be the MSD gene in any other group of vertebrates, its suggested functions of reproduction and fertility (Bircsak et al., 2020; Hansen et al., 2014; Hernández-Ochoa et al., 2009; Ohtake et al., 2008) support the hypothesis that it directly represents, or is tightly associated with the true MSD gene. If *ahr* is indeed the MSD gene in elasmobranchs, its location within the non-recombining region of the X chromosome (Fig. 5) rather than the Y chromosome, would suggest that sex determination is a dosage-dependent sex determination system, as proposed by Wu et al. (2024). Future investigations, including expression data during development and sexual maturation/differentiation, are required to verify *ahr* or identify alternative MSD candidates across diverse shark and ray species.

### PARs, HoxC genes, and zbtb39

Our results also reaffirm the utility of collecting short-read data to map onto whole genome reference sequences for inferences of sex chromosomes. The reassignment of the X chromosome of the thorny skate reference genome to chr46 in this study changes the inference about the evolution and the potential shared ancestry of the elasmobranch X chromosomes (Wu et al., 2024). Based on our X chromosome assignment in this species, available evidence now points to a single evolutionary event for the XX/XY chromosomes of all elasmobranchs.

The application of whole genome short-read sequencing also allowed us to reveal the potential ranges of PARs in the white shark and epaulette shark genomes. Compared to previously discovered PARs in other elasmobranchs, our study indicates variable PAR sizes among species: The white shark has a PAR pair that is ∼ 2Mb long on both X and Y chromosomes, similar to the XPAR of zebra shark (∼ 1.5 Mb, Yamaguchi et al., 2023) and white-spotted bamboo shark (∼ 1.5 Mb, Wu et al., 2024), while the epaulette shark has a PAR that is potentially only ∼300 Kb and the thorny skate has no distinguishable PAR. However, gene annotations on the X and Y chromosomes of the white shark indicate that genes on the ∼2Mb XPAR and ∼2Mb YPAR are mutually exclusive, which suggests a longer, ∼4 Mb PAR is shared between the X and Y chromosomes of white shark, and not currently captured by the original interpretation of the genome assembly. Additionally, short-read mapping results suggest that the XPAR of epaulette shark may be incomplete (Fig. 2; Fig. S3), and the YPAR cannot be unambiguously distinguished. Interpretations of PAR ranges among species, therefore, require caution, as PAR sequences seem prone to fragmentation and misassembly even in genome assemblies with relatively high contiguity metrics (Fig. 1).

Despite potential uncertainties, associations can be found between the *HoxC* clusters and the PARs among the shark species (Elasmobranchii: Selachii) as previously observed in the X chromosomes of the zebra shark (Yamaguchi et al., 2023). In this study, we identified a similar association in the white shark and, potentially, the epaulette shark, and expanded this observation to the Y chromosomes. In the white shark, *HoxC* genes are located within the identified putative YPAR, while in the epaulette shark, *HoxC* genes were identified on scaffolds associated with both the X and Y chromosomes, where the female-to-male coverages were closer to 1 than 2 or 0 (Fig. S3). Searches for the likely boundaries of PARs in the sharpnose sevengill shark (Order Hexanchiformes) and horn shark (Order Heterodontiformes) genomes may, therefore, be guided by the locations of where *HoxC* genes are identified, located at the terminus of the X chromosomes. However, *HoxC* genes have not been identified from the smaller spotted catshark or the white-spotted bamboo shark genomes.

Among the batoids, the presence of *HoxC* genes within this cluster appears to be lineage-specific. While the *HoxC* genes seem absent from the rajiform thorny skate or little skate genomes, they were found in every other examined batoid order in this study. In the torpediniform Caribbean electric ray genome, five *HoxC* genes were identified within the region on the candidate X chromosome (i.e., chr12) that are homologous to the X chromosomes of other elasmobranch species. Likewise, several *HoxC* genes were identified on chrX of the rhinopristiform sawfish and chrX1 of the myliobatiform Atlantic stingray and lesser devil ray. Other than the explanations of incomplete genome assemblies or annotations, the absence of the *HoxC* gametologs on chrY of the two myliobatiform species can potentially indicate the halt of recombination between the PARs occurred after chromosome fusion, or the absence of PAR in Batoidea (given its absence in thorny skate chrX). However, the existence of PARs in most batoid species remains to be examined in future studies.

The locations of the PARs also seem to be associated with a non-recombining *zbtb39* gene on the X and Y chromosomes. In species where boundaries of potential PARs were identified, zbtb39 was found to be closely associated with the PAR. In the more recently assembled white-spotted bamboo shark genome, Wu et al. (2024) identified *zbtb39* to be very close to the PARs on the X and Y chromosomes. In an earlier zebra shark genome assembly (sSteFas1.1; GCA_022316705.1), *zbtb39* was located ∼1.2 Mb upstream of the identified PAR (Yamaguchi et al., 2023) on the X chromosome (annotated as chr41). In the current study, *zbtb39* homologs were found to be ∼1 Mb away from the XPAR and YPAR of the white shark genome, potentially supporting a cross-order association between the locations of *zbtb39* and PARs. However, if the short region on the chrX of the epaulette shark genome indeed represents a PAR, the location of the *zbtb39* homolog would be sitting on the opposite end of this chromosome.

While the presence of *zbtb39* on X chromosomes (X1 for Myliobatiformes) was identified as universal among the examined elasmobranchs, their Y gametologs were found on only three out of four Y chromosomes identified. In the epaulette shark genome, *zbtb39* was missing from chrY and its unlocalized scaffolds despite having relatively high contig N50 among the examined genomes (∼8.8 Mb, Fig. 1, Table 1). The likely *zbtb39* Y gametolog in epaulette shark was found to be on a 153,158 bp long unplaced scaffold, NW_026869144.1, along with another protein-coding gene, zinc transporter ZIP10, whose homolog is found on the X chromosome. Similar situations were likely reflected in the search for EXCGs, where four genes were found to be located on the X chromosomes of most genome assemblies, but on unplaced scaffolds of one genome (Supplementary Table 4). This issue likely lowers the discovery of EXCGs and resonates with the suggestion by Wu et al. (2024) that the suspected male sex-determining factors may lie within the unassembled elements, awaiting detection when complete, error-free, assemblies of the notoriously difficult Y chromosome (e.g., Rhie et al., 2023) become available in the future.

### Speculation: The Preservation of the XY Sex Determination

The long-conserved XY sex determination in sharks and rays is consistent with sex-determining factors emerging on a proto-sex chromosome, subsequently leading to downstream recombination arrest and the degeneration of Y chromosomes. Based on our results, sex chromosome turnover has likely not occurred. This may have been the consequence of a stable environment and low mutation rate, where the mutational load on the non-recombining sex chromosome was largely kept in check.

The consistent presence of *HoxC* cluster genes on the X and Y chromosomes across distantly related elasmobranch species points to long-term purifying selection, albeit to varied degrees (Hara et al., 2018; Yamaguchi et al., 2023), along with the X-Y gene pair recurrently identified in the non-recombining region among these taxa, *zbtb39*. If this non-recombining region is selectively preserved, the chromosomal section with low X-Y sequence divergence (‘young stratum’ in Wu et al., 2024) may represent an ancient region that could be alternatively an older stratum that has been under heavy selective pressure since its recombination arrest across deep time (but see next section).

### Speculation: Did Some Sex Chromosomes Escape Degeneration?

Interestingly, although genome assemblies exhibit sex chromosomes with highly diverged sequences among sharks and rays, cytogenetic studies have not reported heteromorphic sex chromosomes across a large phylogenetic swath of examined species (e.g., Asahida et al., 1995; Maddock & Schwartz, 1996; Schwartz & Maddock, 2002; Uno et al., 2020). This is inconsistent with the canonical model of sex chromosome evolution, where Y chromosomes are expected to degenerate after recombination suppression, leading to heteromorphic sex chromosomes (reviewed in Zhu et al., 2024). In zebra sharks and whale sharks, where both cytogenetic and genomic evidence has recently been examined, no heteromorphic sex chromosomes were identified from the karyotyping efforts (Uno et al., 2020). However, the genomic sequences based on the f-to-m coverage ratio suggest highly differentiated chrX and chrY sequences (Yamaguchi et al., 2023). This discrepancy may indicate young but diverging sex chromosomes or the enlargement of chrY due to the accumulation of heterochromatin or amplicons (reviewed in Schartl et al., 2016). Further investigations on these differentiated sequences can benefit from assemblies of the chrY or cytogenetic techniques such as FISH, as applied in the verification of the bamboo shark chrY in Wu et al. (2024).

Homomorphic sex chromosomes are often associated with lineages whose sex chromosomes are “young” as a consequence of frequent turnover (e.g., Blaser et al., 2013). However, for elasmobranchs, this explanation is inconsistent with the ancient and conserved sex chromosome pairs recently reported (Wu et al., 2024; this study). In our estimation, two candidate mechanisms can potentially explain how the ancient and likely non-recombining Y chromosomes have avoided degeneration.

In the first instance, the Y chromosome may have experienced a slow rate of degeneration as a result of the low mutation rate or a reduced fixation rate of recombination-arresting mutations (Vicoso, 2019). This mechanism has been proposed to partly explain the preservation of W chromosomes in ostriches (Yazdi et al., 2020). Given the extraordinarily slow mutation rate observed in elasmobranchs (Sendell-Price et al., 2023), Y degeneration may occur at slower rates for some species. However, lineage-specific processes have still led to the reduced size and gene content of chrY in other species, such as the epaulette shark and white-spotted bamboo shark, potentially caused by independent occurrences of recombination suppression by inversions or mechanisms acting in gradual manners (Kent et al., 2017).

The second candidate mechanism would be via occasional X and Y chromosome recombination, possibly through sex reversal that results in fertile XY females. This scenario would allow for some recombination between the X and Y chromosomes, potentially breaking up deleterious mutations and creating Y haplotypes with lower mutational loads (Perrin, 2009). This “fountain of youth” mechanism was proposed to explain the lack of differentiation between the XY chromosome pairs in some tree frog (Perrin, 2009; Rodrigues et al., 2018) and seahorse (Long et al., 2023) species, despite apparent recombination suppression in males. Even rare recombination between X and Y could be sufficient to stop the accumulation of mutations leading to XY differentiation (Grossen et al., 2012) or turnover (Blaser et al., 2013).

One consequence of the hypothesized “fountain of youth” mechanism is that sections of sequences on non-recombining chrY should be more similar to the chrX of the same species than they are to the homologous regions on the non-recombing chrY of its sister species (Perrin, 2009). This is consistent with the high similarity observed between the X and Y gametologs of *zbtb39* (Wu et al., 2024), despite the gene being located outside of the PARs in the species examined.

Although sex reversal has not been reported in sharks and rays, many shark species exhibit dynamic sexual traits, with frequent observations of individuals presenting varying degrees of intersexuality, potentially functioning as male, female, or both (Hendon et al., 2013). The rare but non-negligible cases where identification based on observation of the phenotype (presence or absence of claspers) and genetically identified sex conflict (Devloo-Delva et al., 2024; Lee et al., 2024) may also suggest the existence of rare sex-reversed individuals. Additionally, the preservation of Y chromosomes through sex reversal may have occurred in the past, delaying Y degeneration in some species, despite no longer being visible in modern populations.

The direct observation of sex-reversed XY females is likely rare. However, searches for species that potentially exhibit such mechanisms can be guided by the observations of skewed sex ratios in one clutch or litter (e.g., higher male-to-female ratio due to XY females, Matsuba et al., 2008) if genotyping loci on the sex chromosomes or examining their sex-linked markers is available. Caution should also be taken as, even in sex-reversed individuals, recombination between the X and Y chromosomes may not occur. In mice, while recombination rates are determined by the phenotypic sex in XY females, the X and Y chromosomes do not recombine, possibly due to their highly diverged structures (Lynn et al., 2005). Further investigations based on extensive sampling with reliable reporting of morphological sex and the examination of genetic sex through identified sex markers or karyotyping would provide a way to test this hypothesis.

### Speculation: Potential Role of Sex Chromosome Fusions in Myliobatiformes

In this study, we demonstrated that the two mylibatiform species are unique in exhibiting two genomic fusion events that involve the sex chromosomes, which resulted in the formation of a neo-Y chromosome. This is likely shared by other myliobatiform species with multiple sex chromosomes (e.g., Cruz et al., 2011). In the common ancestor of these species, fusion events might have linked potentially sexually antagonistic genes originally occurring on autosomes with sex-determining factors, leading to physiological and/or behavioral changes. As sex chromosomal fusions in elasmobranchs are so far only reported in a subset of Myliobatiformes (along with the possibility in Torpediniformes), the potential incompatibility caused by the fusion might facilitate reproductive isolation (Kitano et al., 2009; Yoshida et al., 2023), although comprehensive investigations involving wider taxon sampling are required. Searches for the functional links and the potential sexual antagonistic effects on chrX1 and chrY would likely yield insights relevant to testing this idea.

Interestingly, a similar fusion event was also inferred in the candidate sex chromosome of the Caribbean electric ray genome (Order Torpediniformes). Both of these events involve the conserved X-linked genes and one identical ancestral autosome. The fusion between the two chromosomes potentially represents a single event that occurred in the common ancestors of Torpediniformes, Rhinopristiformes, and Myliobatiformes, with the secondary loss of fusion in Rhinopristiformes (Fig. 4). Alternatively, the ancestral sex chromosomes and this autosome may possess properties that facilitated the fusions in the two orders who that shared a common ancestor approximately 194.8 million years ago (Stein et al., 2018). However, due to the limited taxa sampled and the uncertain identity of the sex chromosomes in the electric ray genome, we reserve further speculations until additional evidence is available. Future studies on these fusion events may provide valuable insights into the mechanisms of chromosome fusion in elasmobranchs.

### Limitations of the study

In our analyses, we did not include the newer and more comprehensively analyzed bamboo shark reference genome assembled by Wu et al. (2024) as it was not publicly available at the time of our study. This newly assembled genome has improved contiguity relative to the white-spotted bamboo shark genome included in this study, ASM401019v1, labeled as a reference genome of the species. ASM401019v1 has a lower chromosome count (2n = 102) than that reported (2n=106) by Uno et al. (2020), likely due to the high number (101,925) of unplaced scaffolds in the genome complicating the scaffolding processes. The future adoption of the newer white-spotted bamboo shark genome provided by Wu et al. (2024) will likely allow us to discover more syntenic relationships among chromosomes and to discover more EXCGs. That said, we do not expect major changes at the ordinal level because another orectolobiform genome (i.e., the epaulette shark) with excellent contiguity statistics was included in our analyses.

It is also noteworthy that our synteny analyses suggest a more dynamic genomic fusion/fission history among elasmobranchs than past work, as shown by the gene mapping results between the closely related whale shark and zebra shark reported by Yamaguchi et al. (2023), but higher than implied by Wu et al., (2024). In our analyses, the pairwise syntenic relationships established by our 1-to-1 ortholog analyses are generally supported by the whole genome alignment based on Progressive Cactus (Armstrong et al., 2020) and halSynteny (Krasheninnikova et al., 2020) (Supplementary Fig. 5) in species where both types of data are available. Due to the different sets of samples and methodologies, we suspect that the stringent and conservative methods applied by Wu et al. (2024) may have resulted in the discovery of a different set of orthologous pairs than the ones we present herein. Reconciling the two sets of results in the future should bring a richer understanding of the evolutionary dynamics associated with synteny.

Finally, we acknowledge that the inferences made in this study are limited by the completeness of genome assemblies and the fact that four out of the 13 elasmobranch orders are not represented in this study. Elasmobranch species are known to exhibit diverse physiological systems, such as in their reproductive mechanisms (Nakaya et al., 2020), multiple cases of facultative parthenogenesis (e.g., smooth-hound shark, Esposito et al., 2024), XX/XO system (*Potamotrygon* sp. C, De Souza Valentim et al., 2013), and a species with a possible, but not confirmed, ZZ/ZW system (*Hypanus americanus*, Maddock & Schwartz, 1996; Schwartz & Maddock, 2002). Our current understanding of elasmobranch karyotype evolution still leaves open the possibility for the discovery of alternative and independently evolved sex determination systems. With the efforts for generating high-quality genome assemblies such as the VGP and the Squalomix project (https://github.com/Squalomix/info/), we can confidently expect additional insights into elasmobranch sex determination to come to fruition in the near future.

Our study was conducted independently but in parallel to a study by Niwa et al. in preparation, although neither group had prior knowledge of each other’s efforts until recently. We encourage readers to refer to Niwa et al.’s article for an alternative perspective on elasmobranch sex chromosome evolution. While the articles have different focuses, they are complementary to one another.

## Conclusions

In summary, our study revealed a more ancient origin of the shared elasmobranch sex chromosome pair than inferred previously (∼265.1 Mya, Stein et al., 2018). This is considerably older than the shared XY system documented for therian mammals (∼166 – 190 mya, Graves, 2016) and the sturgeon ZW system (∼180mya, Kuhl et al., 2021). Our comparative synteny analyses suggest that there have been two fusion events associated with the sex chromosomes of Myliobatiformes. The data suggest there was a fusion between the ancestral XY and two autosome pairs, resulting in two X chromosomes and a neo-Y chromosome observed in the two species examined. Although our analyses do not reveal any master sex-determining genes on the Y chromosome, *ahr-like* and other EXCGs on the X chromosomes provide promising candidates for future investigations of a shared ancient dosage-based system, such as the ZW system in the birds (Ioannidis et al., 2021).

## Acknowledgments

We thank Linelle Abueg and Tatiana Tilley for their contributions to the generation of the genome assemblies. We thank the Centro Nacional de Analisis Genomico (Barcelona, Spain) for the generation and curation of the lesser devil ray genome. We thank Bianca Prohaska, Venkatesh Byrappa, Stacia White, Dean Grubbs, Kady Lyons, and Chris Lowe for their contributions to acquiring samples used for genome assembly, and Manfred Schartl, Mark Erdmann, Frank Tulenko, Arve Lynghammer, Jeff Kneebone, and Sabine Wintner for their contributions to the obtaining the samples or sequences in the short-read sequencing dataset.

## Author contributions

S.-H.L. and G.J.P.N. conceived the study; G.J.P.N. and J.M. coordinated the samples; O.F., E.J., J.B., N.J., T.T., B.O, B.H., and A.R. performed sequencing and genome assembly; O.F. and J. B. generated the genome assembly methods and descriptions; N.B., D.A., S.P., D.-L.P., J.M.D.W., and K.H. curated the genomes; P. L. examined the thorny skate genome; L.S-.C., E.H., and D.M. provided the genome-wide sequence alignment data; S.-H.L. performed the sex chromosome-related analyses and interpreted the data; S.-H.L. drafted the manuscript; E.H., D.M., and G.J.P.N. participated in revising the manuscript.

---

**Supplementary Figure 1.**
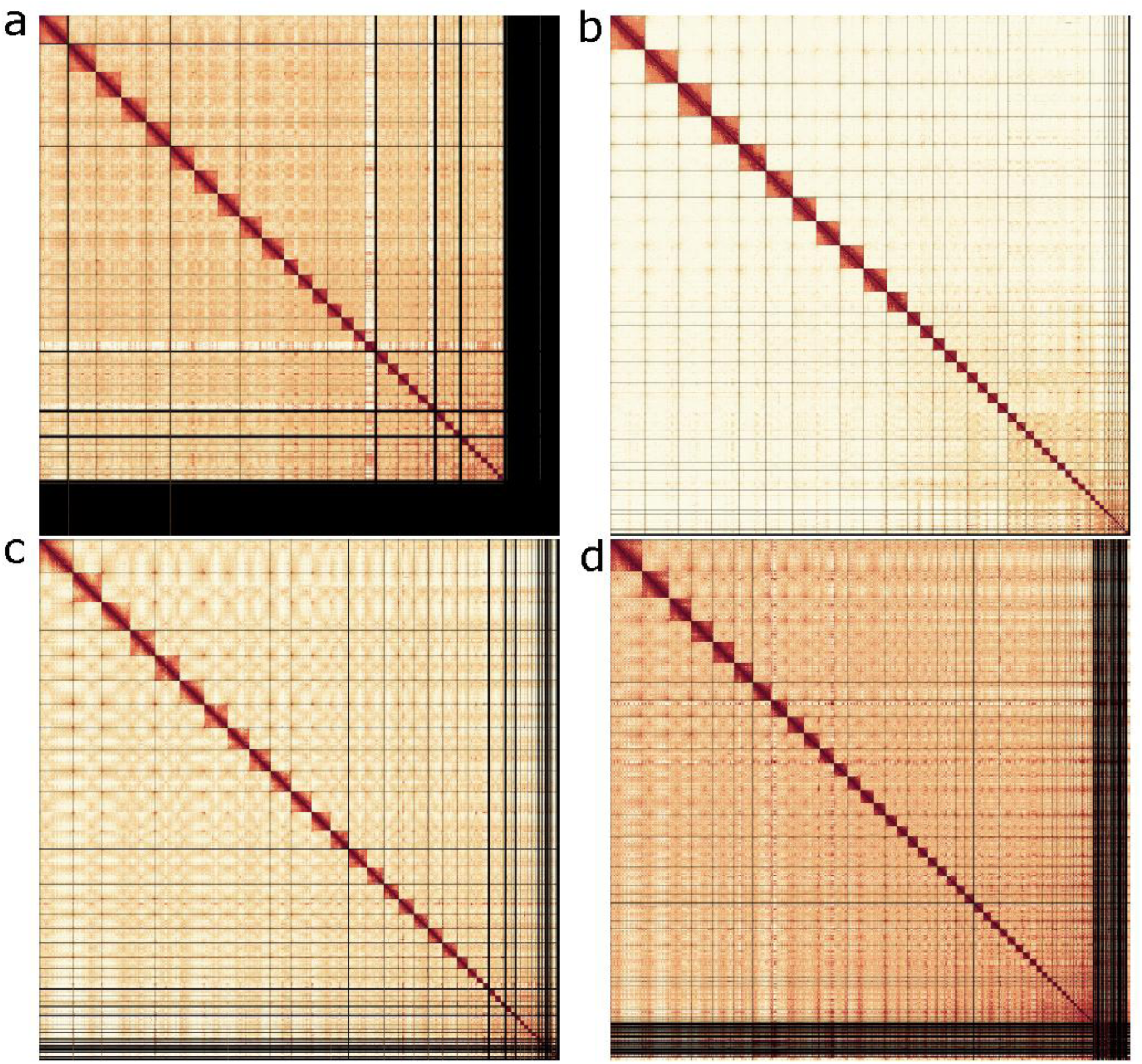
Hi-C interaction heat maps for the four newly assembled elasmobranch genomes after curation, visualized with PretextView (https://github.com/sanger-tol/PretextView). (a) Atlantic stingray. (b) Smalltooth sawfish. (c) White shark. (d) Zebra shark.

**Supplementary Figure 2.**
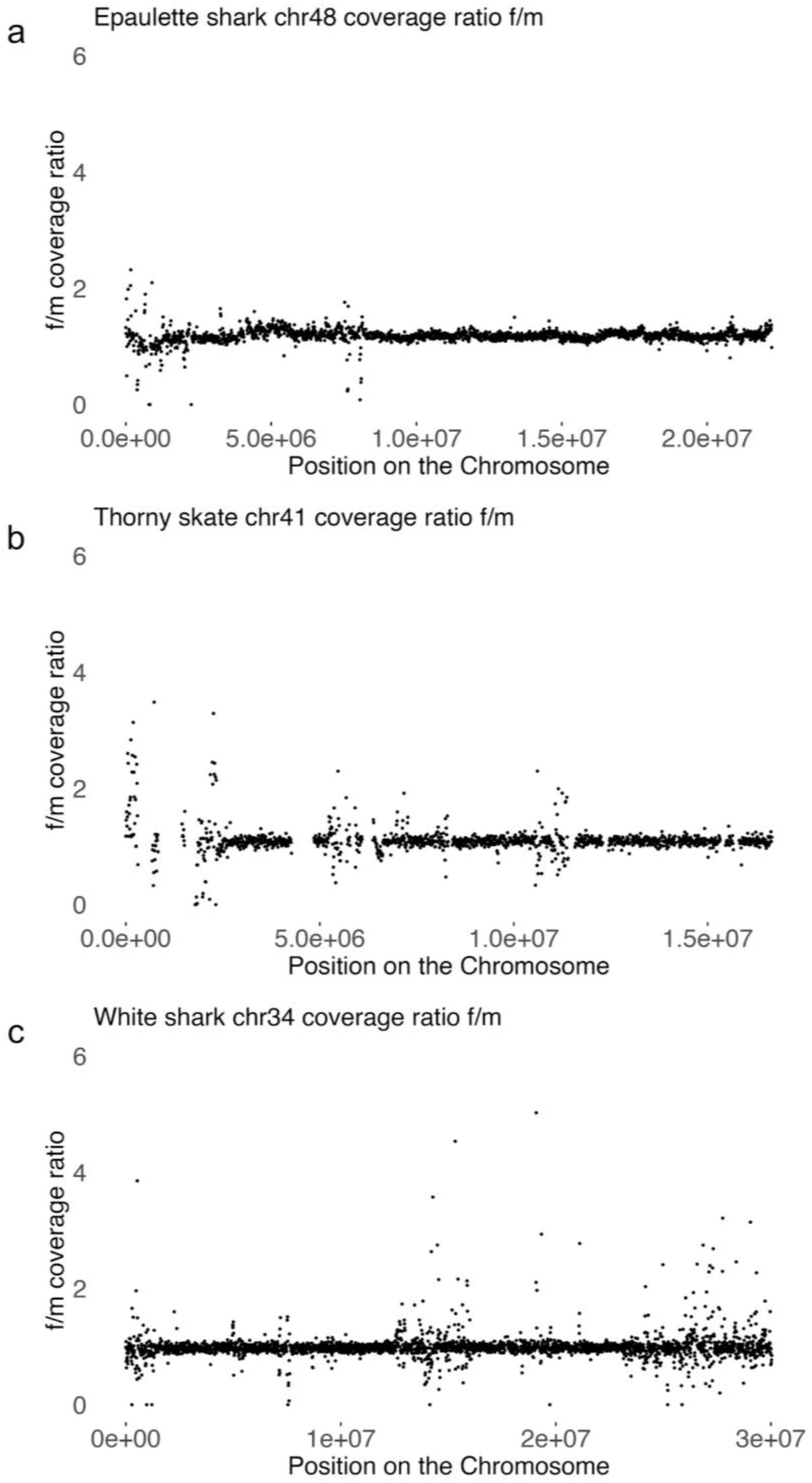
Pooled female-to-male read coverage ratio across one autosome in each species based on the mapping of short-read sequencing data to the reference genome assemblies. Ratios along the chromosomes are visualized in 10,000 bp, non-overlapping windows. Chromosomes are selected based on their similar size to the annotated X chromosomes. (a) chr48 of epaulette shark (b) chr41 of thorny skate (c) chr34 of white shark.

**Supplementary Figure 3.**
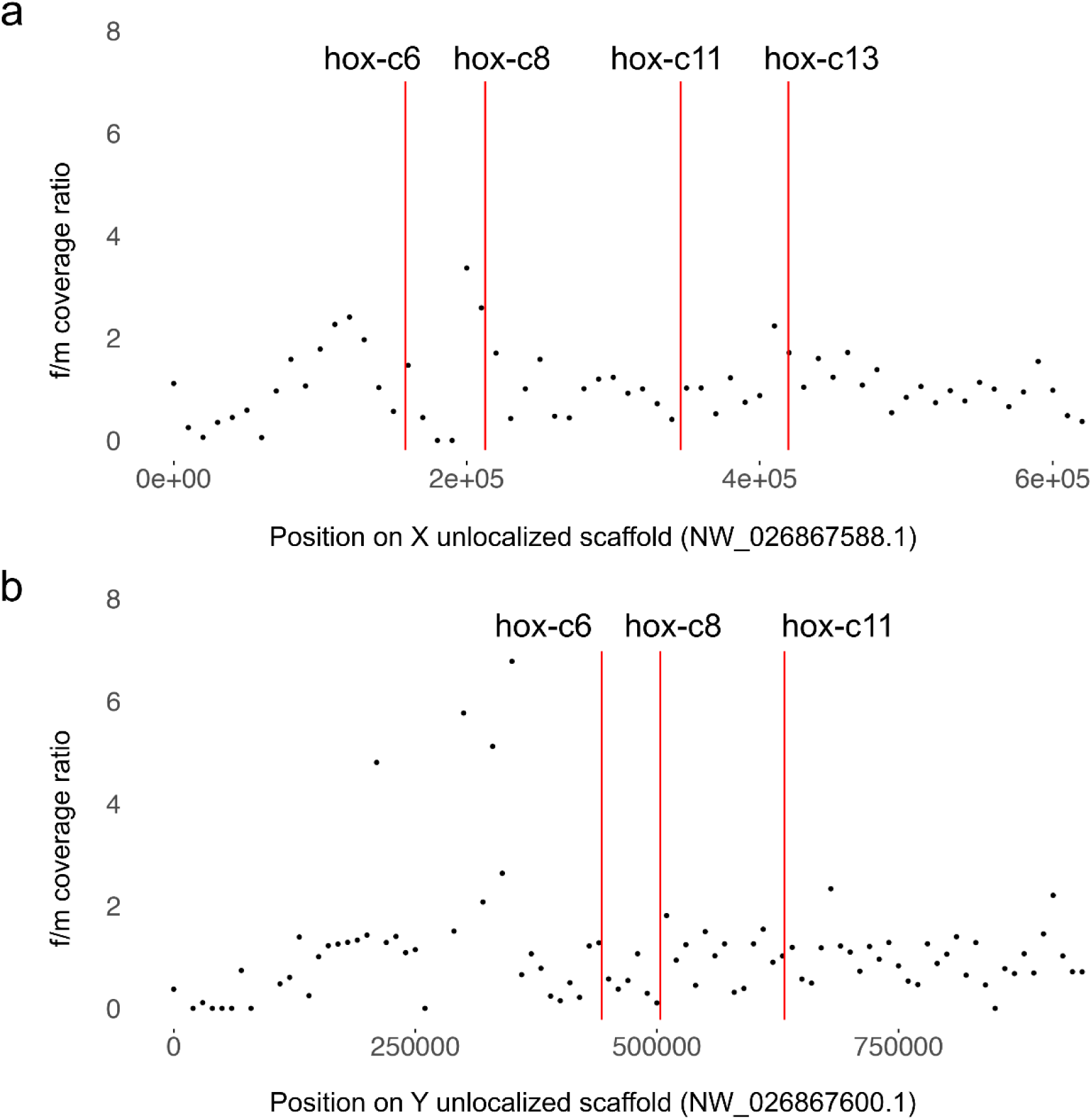
Pooled female-to-male read coverage ratio across (a) the X unlocalized scaffold and (b) the Y unlocalized scaffold where *HoxC* genes are identified. The approximate locations of *HoxC* genes are labeled.

**Supplementary Figure 4.**
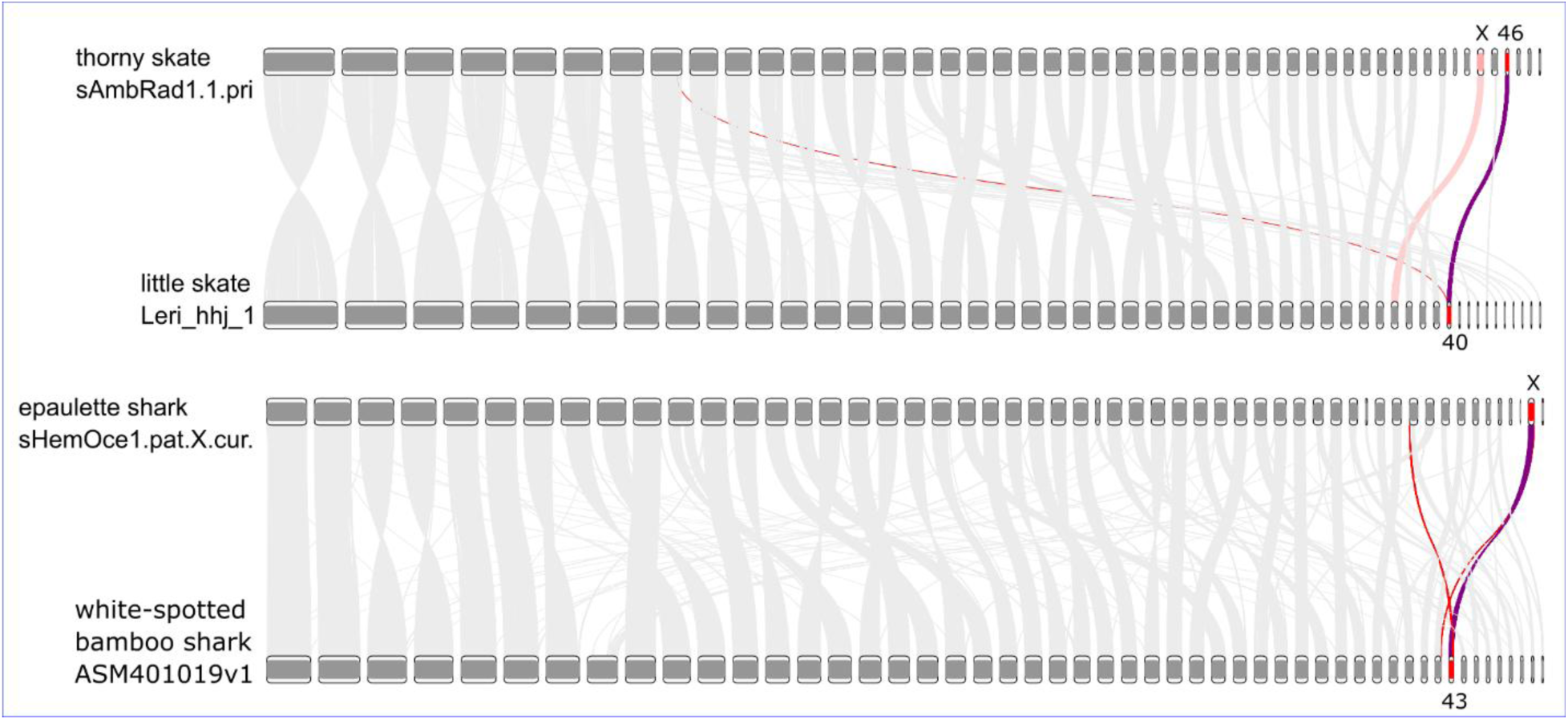
Synteny between the thorny skate and little skate reference genome assemblies (top) and between the epaulette shark and white spotted bamboo shark reference genome assemblies (bottom). Chr40 of the little skate genome and Chr43 of the white-spotted bamboo shark genome (see Discussion) are inferred as ChrX of the species. Annotated or inferred chrXs are highlighted in red, the homology paths that indicate shared relationships are highlighted in purple, while paths that have one end to a chrX are highlighted in red.

**Supplementary Figure 5.**
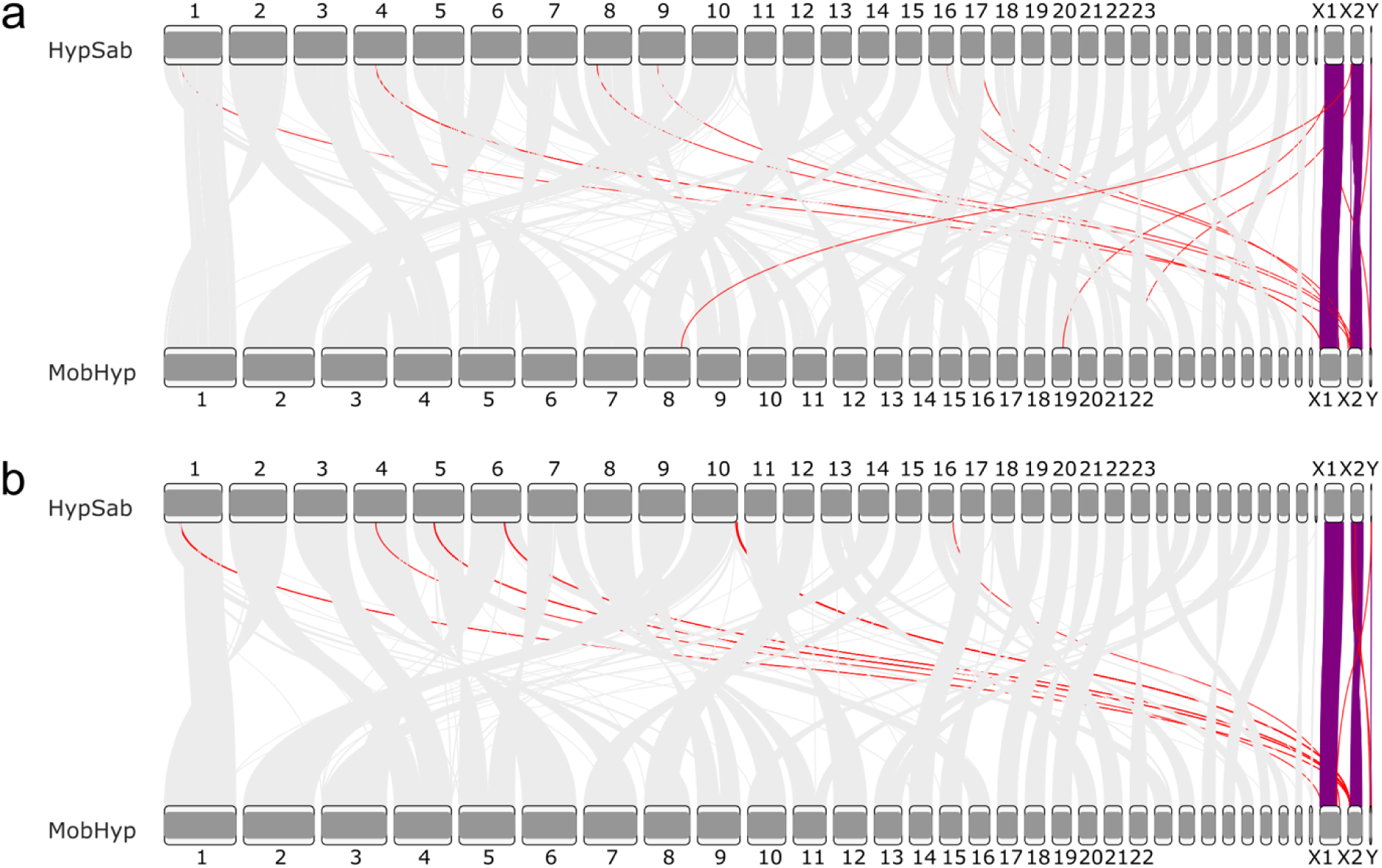
Syntenic relationships between the Atlantic stingray and lesser devil ray reference genome assemblies inferred from (a) 1-to-1 ortholog analyses and from (b) progressive cactus alignment and halSyntenty. Annotated or inferred chrXs are highlighted in red, the homology paths that indicate shared relationships are highlighted in purple, while paths that have one end to a chrX are highlighted in red.

